# High-intensity sheep grazing impoverishes soil seed banks in sand grasslands

**DOI:** 10.64898/2026.03.18.712656

**Authors:** Gergely Kovacsics-Vári, Judit Sonkoly, Katalin Szél-Tóth, Andrea McIntosh-Buday, Luis Roberto Guallichico Suntaxi, Szilvia Madar, Patricia Elisabeth Díaz Cando, Viktória Törő-Szijgyártó, Béla Tóthmérész, Péter Török

**Author notes:** Corresponding author Péter Török, Department of Ecology, University of Debrecen, Debrecen, Hungary.

## Abstract

The effects of the selection of livestock type (e.g., sheep or cattle) and grazing intensity on the soil seed bank of sand grasslands of conservation interest were studied. 25 grazed grassland sites classified into four grazing intensity categories were studied. The soil seed bank was analysed by seedling emergence; germinated seedlings were classified into morpho-functional, social behaviour type (SBT) and CSR strategy groups. The following hypotheses were tested: i) Diversity and density of soil seed banks are lower in sheep-grazed sites than in cattle-grazed ones. ii) The species composition, diversity, and density of the soil seed banks are more strongly affected by grazing intensity than by the livestock type. iii) Leaf traits, SBT and CSR strategy composition are highly affected both by livestock type and grazing intensity. The main effect of livestock type only affected seed bank density, while that of grazing intensity had a significant effect on most of the variables. Most of the studied variables were affected by the interaction of grazing intensity and livestock type. Total seed bank density was lower at all grazing intensity levels in sheep-grazed sites than in cattle-grazed ones, especially close to frequently visited places. We found that sheep grazing sustained a much lower total seed bank density and lower density of species of natural and semi-natural habitats regardless to the grazing intensity. Thus, livestock type must be carefully selected and high-intensity sheep grazing should be avoided in the long-run when managing sand grasslands.

**Highlights:**

- The soil seed banks of sheep and cattle grazed sand grassland were studied
- Effect of grazing intensity found the most important driver of seed bank diversity and density
- The total soil seed bank density was higher in cattle than sheep grazed sites
- Both intensity and livestock type must be considered in the grassland management planning
- High intensity sheep grazing should be avoided in sand grassland management

## 1. Introduction

Natural and semi-natural grasslands globally have high conservation value by acting as refuges for many animal and plant species (Didukh, 2009). They also provide important ecosystem services including food supply for people and their livestock (Squires et al., 2018). Similar to other ecosystems, natural and semi-natural grasslands are facing a series of conservational problems. Besides area loss, the most widespread problem is the change of management type and -intensity in grasslands, ranging from complete abandonment to overuse in the form of high -intensity mowing and/or livestock grazing (Török & Dengler 2018).

From a conservation perspective, grazing by domestic livestock is considered to mimic wild grazers and thus it is frequently applied in sustainable grassland management (Senn, 2022). It was stressed by several authors that the effect of grazing on the subjected vegetation is strongly dependent on grazing livestock type, grazing intensity, and the productivity and diversity of the subjected grassland (Rook et al., 2004; Fraser et al., 2022; Török et al., 2024). Thus, while grazing is extensively studied globally, it is difficult to identify general patterns or form universal recommendations for the sustainable use of grasslands for many regions and there are still numerous open questions waiting for answers (Fraser et al., 2022; Jiang & Wang, 2022). For example, belowground aspects including the diversity and density of soil seed banks are less considered than aboveground vegetation (but see Dreber & Esler, 2011; Chu et al., 2019; Li et al., 2024).

The role of livestock type in understanding the effects of grazing is less studied than that of grazing intensity, however, it is just as important as it affects the biodiversity and sustainability of grasslands (Fraser et al., 2022; Jiang & Wang, 2022; Török et al., 2024). There are several studies comparing the effects of sheep and cattle grazing (e.g., Tóth et al., 2016; Kovacsics-Vári et al., 2023; Rodriguez et al., 2023). In the vegetation of alkaline grasslands and alpine meadows, Tóth et al. (2016) and Rodriguez et al. (2023), respectively, found that livestock type (cattle or sheep) is a more crucial driver of vegetation composition than grazing intensity. However, the opposite was found in sand grassland vegetation by Kovacsics-Vári et al. (2023). These findings raised the question of what effect the livestock type and grazing intensity can have on the composition and density of the soil seed bank (SSB) of sand grasslands. Because the grazing selectivity and behaviour of sheep and cattle differ, they presumably have different effects on vegetation composition. For example, sheep consume plants closer to the ground and prefer vegetative parts, while cattle prefer reproductive parts but are more selective for vegetation patches with higher biomass (Metera et al., 2010; Jerrentrup et al., 2015). In the long run, the difference in grazing behaviour can also affect the species composition and diversity of the SSBs. Furthermore, Tóth et al. (2016) found that the vegetation of cattle-grazed sites has a higher functional and taxonomic diversity compared to sheep-grazed ones. However, these expectations were not confirmed by a former comparison of sheep and cattle grazing in sand grassland vegetation as livestock type was found to be a less significant driver of the selected variables (Kovacsics-Vári et al., 2023). We presumed that these differences may affect SSB characteristics too (Shi et al., 2022). Based on the previous findings, we expected that grazing intensity is a more crucial driver of the SSB than livestock type in sand grasslands.

Grazing has been shown to affect the functional trait and plant functional type composition of grasslands (e.g. Díaz et al. 2007; Papanikolaou et al., 2011; Jiang et al., 2023), which has also been documented in sand grasslands (Kovacsics-Vári et al., 2023). Besides plant functional types, the CSR strategy system, originally formulated by Grime (1974) and reconsidered for example by Hodgson et al. (1999) and Pierce et al. (2013), offers a simple and comprehensive system for the characterisation of plant species based on their ecological strategy. Despite being simple and comprehensive, CSR strategies are not regularly applied for studying the effects of grazing management (but see e.g., Moog et al., 2005; Teuber et al., 2013). However, the CSR strategy system has a widely accepted relevance, therefore its applicability in grazing management is worth studying. The CSR strategy systems has also been adapted to Central European vegetation and the Social Behaviour Type (SBT) classification of Borhidi (1995) has also been developed based on it, by subdividing the ruderal category into several subcategories. The assignment of CSR strategies reconsidered by Pierce et al. (2013) is based exclusively on leaf traits. The previously mentioned differences in the grazing behaviour of different livestock types (Metera et al., 2010; Jerrentrup et al., 2015) may also affect leaf traits. For example, leaf dry matter content decreases while specific leaf area increases with increasing grazing pressure (Golodets et al., 2009; Cruz et al., 2010; Kovacsics-Vári et al., 2023). Therefore, we presumed that the SSB also reflects the differences in the CSR and SBT strategy type composition of the aboveground vegetation created by the direct influence of grazing on aboveground plant traits including also leaf traits.

SSBs can play a crucial role in community resilience and they are considered as ecological memory acting as a back-up of the community bridging unfavourable conditions (Perkins et al. 2019, Török et al., 2020). While studies dealing with grazing effects on the composition and diversity of vegetation are numerous (e.g. Díaz et al., 2007, Metera et al., 2010), SSBs received much less attention in this respect. Grazing might affect the SSB in many ways including the followings: i) by the selective grazing of reproductive structures grazing decreases the local input of seeds (Wentao et al., 2023), ii) via ecto-and endozoochorous dispersal it can transport propagules in and into the subjected communities (Matus et al., 2005, Ozinga et al., 2009, Eichberg et al., 2007), iii) by their trampling, grazers facilitate the incorporation of seeds into the soil (Eichberg & Donath, 2018). As livestock grazing is increasingly used as a conservation and restoration measure in many regions and habitat types, studying how it affects community resilience in the form of SSBs is also important from a conservation point of view. Former studies on the SSB of sand grasslands analysed its density and species composition and how it changes in response to some management extremes (e.g., overgrazing and abandonment in sand grasslands, Matus et al., 2005). The role of SSBs in secondary succession (Halassy, 2001; Török et al., 2009; Török et al., 2018a) and the functional role of SSBs in the restoration of sand grassland habitats (Matus et al., 2003; Bossuyt et al. 2007, Godefroid et al., 2018) have also been studied. To our knowledge, no study has analysed the composition and density of SSBs in the context of both livestock type and grazing intensity.

Based on the above considerations and the identified knowledge gap, we aimed to study the soil seed bank (SSB) in sheep- or cattle-grazed grasslands used as pastures subjected to multiple levels of grazing intensity. The following hypotheses were tested: i) Diversity and density of SSBs are lower in sheep-grazed sites than in cattle-grazed ones likely due to the higher selectivity of sheep for forbs. ii) The species composition, diversity, and density of the SSBs are more strongly affected by grazing intensity than by livestock type. iii) The leaf trait and CSR and SBT strategy composition of the SSBs are highly affected both by livestock type and grazing intensity.

## 2. Materials and methods

### 2.1. Study area

The study area is located in the Nyírség region, East Hungary, Central Europe, where we have selected 25 sand grassland sites for sampling (Fig. 1). Sand grasslands of the Nyírség are included as "Pannonian and Pontic sandy steppes" (E1.1a) habitat types into the EU Habitats Directive and are critically endangered habitats in Europe (EC Directorate-General for Environment et al., 2017). In general, sand grasslands provide refugia for numerous endemic plant and animal species (Didukh, 2009; Török et al., 2018b). In sand grasslands, i) there is no salt accumulation (in contrast to saline grasslands), ii) they are relatively easy to be cultivated (in contrast to rocky grasslands), and because iii) these grasslands are characterised by a rather open vegetation coverage, they are highly susceptible to the spread of invasive species from closely located man-made habitats (Botta-Dukát, 2008). The Nyírség is characterised by a moderately continental climate. The mean annual precipitation ranges between 530 and 680 mm (Dövényi, 2010), and in arid years it can be even lower than 400 mm (Négyesi, 2018). Mean annual temperature falls between 9.4 and 9.8 °C, the hottest month is July while the coldest is January (mean temperature in Nyíregyháza is 22.7°C and -0.8°C, respectively, based on the 1991-2021 period, Climate-Data, 2025). The mean annual temperature in the year of the vegetation sampling (2021) was 10.8 °C, 11.8 °C in 2022, and 12.3°C in the year of the SSB sampling (2023). On average, 510 mm precipitation was received in 2021, 500 mm in 2022, while with 770 mm this figure was higher in 2023 (Hungarian Meteorological Service, 2025). Typical soil types of the selected sites are Dystric to Brunic arenosols. The soil pH ranges between 4.45 and 5.71, while humus contents between 0.6 and 2.6 m/m% (Kovacsics-Vári et al., 2024).

**Figure 1.**
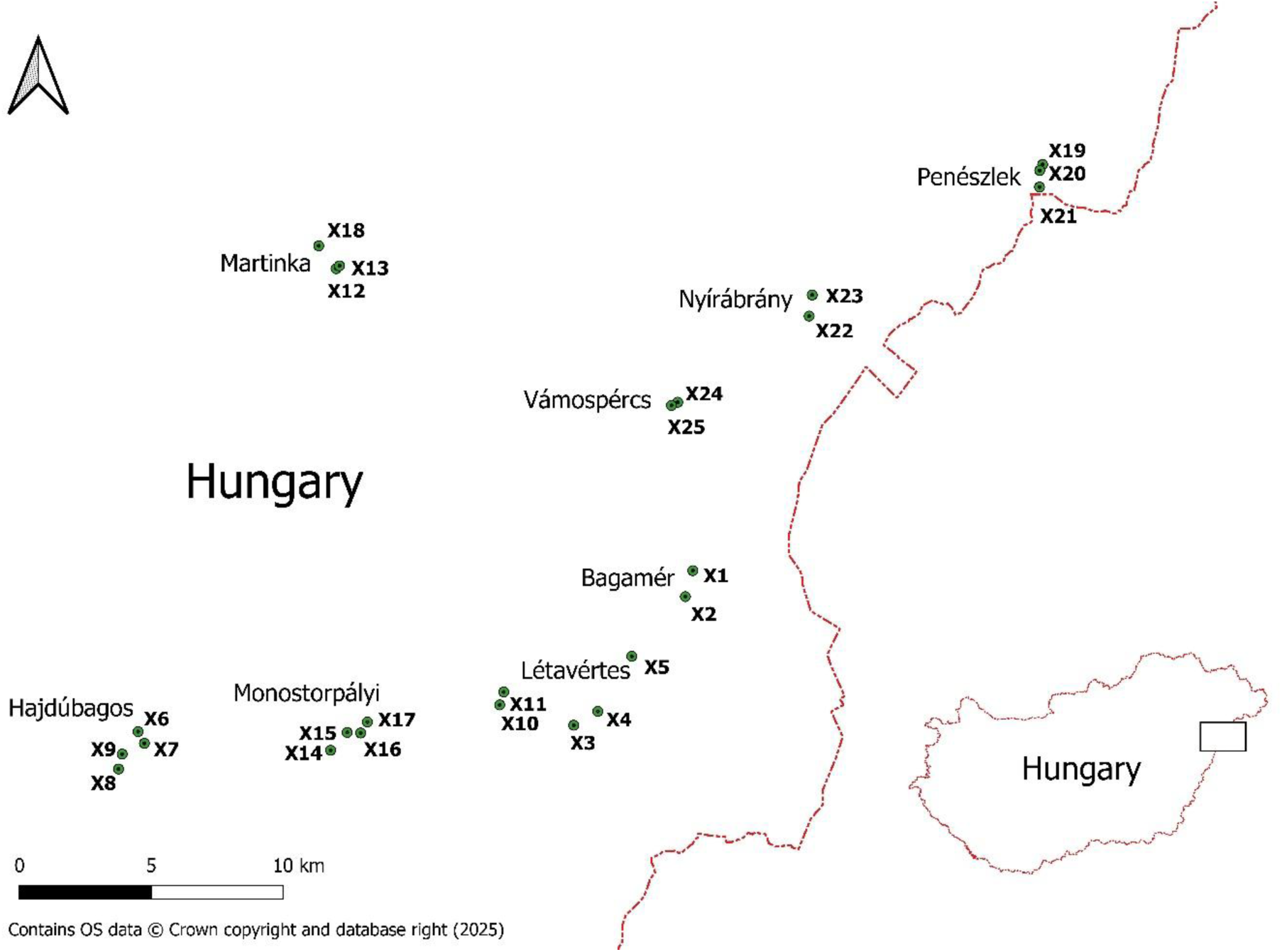
Location of sampling sites in the Nyírség region, Hungary (Central Europe). For detailed information on the grazing intensities and livestock type of sites please see Appendix A.

The Nyírség region is densely populated and tree plantations as well as croplands dominate the region. According to the national park rangers, the effect of wild mammals (e.g., hare or roe deer) on the vegetation is negligible. Our study sites belong to the Hortobágy National Park Directorate and herders lease the area from the directorate; hence they must follow strict management regulations except for one study site, which is owned by the local government. The selected sites are characterized by seasonal, pastoral cattle and sheep grazing and their size ranges between 10 ha and 200 ha. Livestock breeds are Merino for sheep, while Limousin, Hungarian Simmental and Charolais for cattle. The grazing livestock type and intensity of grazing did not change noticeably during the study period. The selected pastures were typically commons belonging to a given settlement and already served this function in the 19th century (Tímár & Biszak, 2010). The elevation above sea level ranges between 102 m and 144 m.

### 2.2. Vegetation composition of the studied grasslands

In the studied grasslands, characteristic perennial graminoid species are *Festuca pseudovina, F. vaginata,* and *Corynephorus canescens*. In the studied grassland types, where livestock grazing is frequent and rather high in intensity, *Cynodon dactylon, Bromus hordeaceus, Anthemis ruthenica*, and *Euphorbia cyparissias* also become typical (Borhidi, 2012). The most typical short-lived forbs of these acidic sand grasslands are *Cerastium semidecandrum, Filago minima,* and *Myosotis stricta* (Borhidi, 2012; Jentsch & Beyschlag, 2003). Furthermore, other perennial forb species sensitive to disturbance such as *Pulsatilla flavescens,* or *Anacamptis morio* can also be found in the most valuable stands of these grasslands (Borhidi, 2012). In grazed sites, some spiny and/or highly unpalatable forbs, e.g., *Verbascum phlomoides, V. densiflorum*, and *Eryngium campestre* can reach relatively high cover. Native weeds including *Chenopodium album, Polygonum aviculare* and some aliens like *Ambrosia artemisiifolia, Conyza canadensis, Setaria viridis,* or *Digitaria sanguinalis* can also be detected (Appendix A).

### 2.3. Experimental design

We selected 25 grazed sand grassland sites in the municipality of eight settlements (Fig. 1). The 25 sites were classified into four grazing intensity categories based on information on grazing intensity provided by national park rangers. We classified the grazed sites first into two main categories based on livestock units/ha data (LU/ha). Then we further refined the categories by considering the distance of the sites from frequently visited points (e.g., watering places). We counted the number of animal droppings to confirm the presence of livestock (see classification in Table 1). In each selected site, we designated 10 × 10 m quadrats. We selected five 4-m^2^-sized plots arranged in a regular pattern within the 10 × 10 m quadrats to obtain a more uniform heterogeneity of the samples in the sampling sites (see sampling design in Fig. 2). We selected 10 × 10 m quadrats closer and further away from frequently visited places. In selecting quadrats, we considered the followings: i) Verges of dirt roads and ditch banks should be avoided. Based on the visual detection of the vegetation patches, we deemed 10 m as a minimum distance from the verges. ii) We avoided sites where large bare ground patches were created by activities other than grazing. iii) At the small scale, we did not sample patches covered by dung of cattle to avoid bias in SSB data. We had altogether 13 sampling sites grazed by cattle and 12 grazed by sheep, 11 grazed by a higher LU/ha, 14 grazed by a lower LU/ha, 12 sampling sites being further away from the frequently visited places while 13 being closer to the frequently visited places.

**Figure 2.**
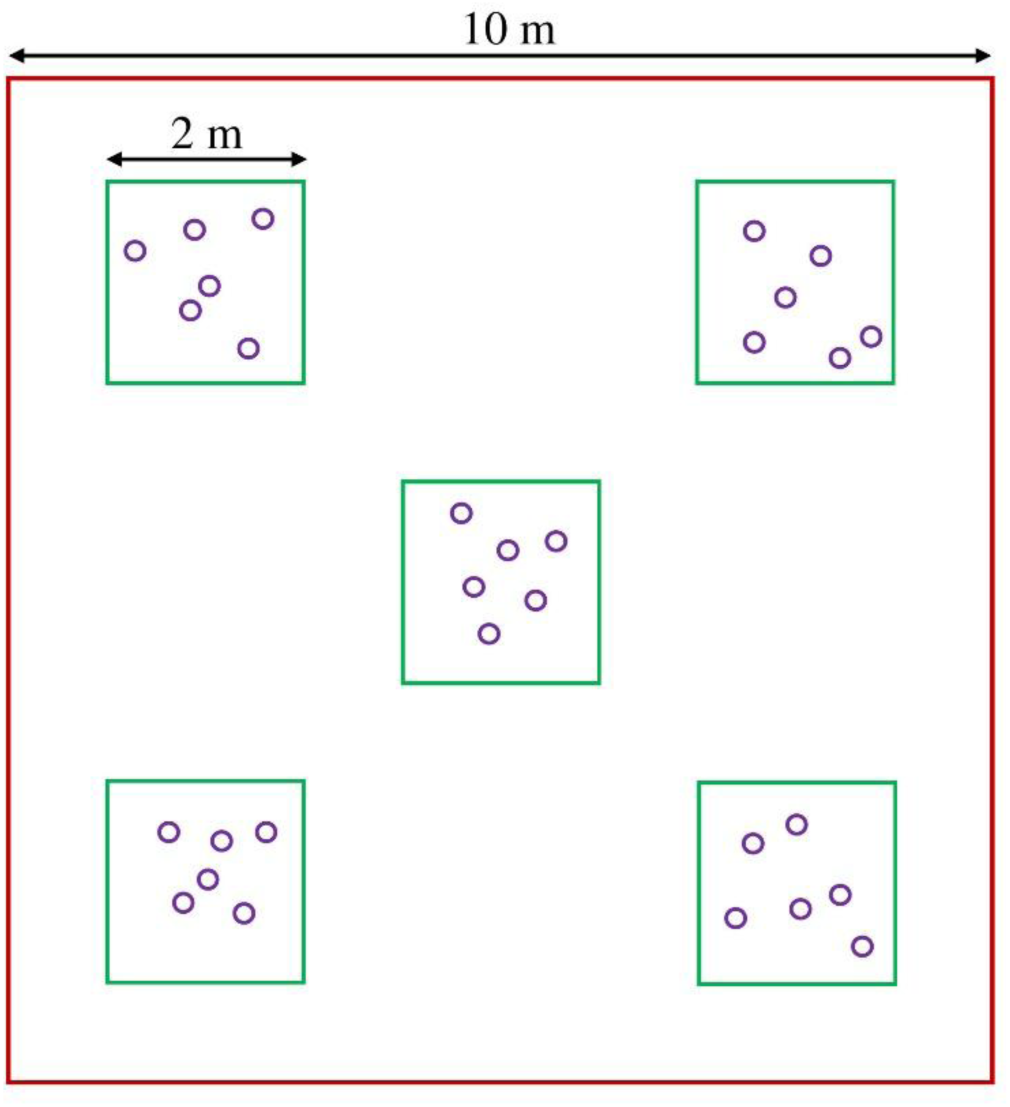
Design of the sampling. Six soil cores (violet circles) were randomly taken from a 4-m^2^-sized plot, and pooled into one sample, hence we had five pooled samples in a sampling site. One core has a diameter of 4 cm, 10 cm depth, and approximately 126 cm^3^ in volume. We assessed the number of animal droppings within the 10 ×10 m quadrats.

**Table 1.** Detailed description of the assignment of grazing intensity levels. Areas located within approximately 150 m of frequently visited places were used by the animals at similar frequencies during the resting period. At a distance higher than 150 m, the animals are herded to different directions, thus the visiting frequency/grazing intensity becomes less high. Notations: LU/ha = livestock unit/hectare. One head of sheep is considered as 0.2 LU, while one head of cattle as 1 LU; FVP = Frequently Visited Place (watering point, resting point, barn).

| Category of grazing intensity | LU/ha | Distance to FVP | Number of droppings |
| --- | --- | --- | --- |
| First | 0.5 – 0.8 | more than 150 m | Low |
| Second | 0.5 – 0.8 | 30 to 150 m | High |
| Third | 1.1 – 4 | more than 150 m | Low |
| Fourth | 1.1 – 4 | 30 to 150 m | High |

### 2.4. Sampling and management of the samples

In the selected 4-m^2^-sized plots were recorded the percentage cover of vascular plant species and collected SSB samples. Recording of percentage cover data was implemented in 2021 from late May to early June, at the peak of the vegetation period. Soil samples for SSB analyses were collected in the spring, early March 2023 using core samplers. We had altogether 125 vegetation plots and also collected 125 pooled SSB samples from the plots in 25 sites with five replications each (Fig. 2). For SSB, six cores (4 cm diameter, 10 cm depth, 126 cm^3^ of volume) originating from the same plot were pooled for germination and analyses. We used the seedling emergence method by Ter Heerdt et al. (1996). Samples were concentrated by sieving, then spread in a thin layer on the surface of steam-sterilised potting soil. The pots containing the samples were placed in a greenhouse without heating. We also used three pots without a sample layer as control, to detect airborne species not being part of the SSB samples. The germination period lasted from early April to mid-November with two sub-periods (a spring period from early April to the end of June and an autumn period from the end of August to mid-November with the inclusion of a mid-summer germination break). Seedlings were regularly counted, identified and then removed from the pots. Unidentified individuals were transplanted to separate pots for later identification. Seedlings and transplanted individuals were regularly checked for appropriateness of soil conditions, detection of decaying plants caused by fungus infections or damage created by molluscs or insects. The frequency of regular checking was higher during the hot weeks to avoid the expected higher mortality by careful watering and manual ventilation of the greenhouse. To mimic summer drought, when no further seedlings emerged, watering of the soil samples was stopped, and only the transplanted individuals were watered between early July and the end of August. The watering break of SSB samples coincided with the typical summer drought in the region and likely enhanced the germination vigour of species requiring a heat-stratification (Baskin & Baskin, 1998). Before the watering break, all seedlings were removed or transplanted from the pots. After mid-November, conditions are not suitable for further germination (low temperatures and short daylight period), therefore we stopped the germination and prepared the transplanted individuals for overwintering. Seedlings were identified based on the nearly 20 years of experience of the senior author, and the identification was supported by a photo collection of seedlings at the Department of Ecology, University of Debrecen. We also used the reference works of Csapody (1968), and Muller (1978) for the identification of seedlings. Nomenclature follows Euro+Med PlantBase (Euro+Med, 2006).

### 2.5. Data collection and processing

Several species groups were pooled during the data processing. *Equisetum* species (*E. arvense*, *E. ramosissimum*) were omitted from the comparison of vegetation and seed bank data as these species does not have seeds. We were not able to differentiate small *Filago* individuals in the field, so *F. minima* and *F. arvensis* were pooled in the analyses, the same hold for *Geranium molle* and *G. pusillum* and for *Polygonum arenarium* and *P. aviculare*. We were not able to grow some seedlings of *Teucrium* species due to their high mortality, so we pooled them as *T. chamaedrys/scordium*. It was also not possible to grow seedlings of *Viola arvensis* and *V. kitaibeliana* until flowering, so we pooled them for the analyses.

We classified the identified plant species of the SSB into four morpho-functional groups based on life forms and morphological characteristics provided by Király (2009) and Sonkoly et al. (2023). These were perennial graminoids, perennial forbs, short-lived graminoids, and short-lived forbs. Annual and biennial species were considered as ‘short-lived’, while all other long-lived species were considered as ‘perennial’. We classified the detected plant species into mixed CSR strategy types following the method and MS Excel tool (StrateFy) of Pierce et al. (2013), where the classification was based on three leaf traits, namely leaf area (LA), specific leaf area (SLA), and leaf dry matter content (LDMC) (Liu et al., 2025). We obtained leaf trait data from the Pannonian Database of Plant Traits (PADAPT, Sonkoly et al., 2023). We classified the detected plant species into Social Behaviour Types (Borhidi, 1995) to represent disturbance tolerance. Borhidi’s (1995) classification is a modified and extended version of the classical CSR strategy types along which we were able to classify the detected species into three species groups with increasing ruderality: i) species of natural and semi-natural habitats: natural pioneers (NP), generalists (G), specialists (S) and natural competitors (C); ii) disturbance tolerators (DT); and iii) ruderal and weedy species: weeds (W), adventives (A), ruderal competitors (RC) and adventive competitors (AC).

### 2.6. Data analyses

We compared the composition of vegetation and seed banks using a PCA ordination visualised in CANOCO 5.0. Livestock type (categorical variable) and levels of grazing intensity (ordinal variable) as independent variables (fixed factors) and site identities (random factor) were included for calculating generalized linear mixed models (GLMM, Poisson probability distribution, loglink function). To improve normality of count data, logarithmic transformation of data (log (x+1)) was used integrated into loglink function of SPSS. Dependent variables of the GLMMs are summarized in Tab. 2. For dependent variables such as leaf traits and CSR strategies, community-weighted means (CWMs, weighted by seed numbers) were calculated and tested. All univariate statistics were calculated using SPSS 26.0 program package (IBM Corp, 2019). The Bonferroni-Holm method (Holm, 1979) was applied to control family-wise error rate by adjusting the raw *p* values within each family of related response variables. For the Bonferroni-Holm correction we used the R environment (version 4.3.2) (R Core Team, 2023).

**Table 2.** Effect of grazing intensity, livestock type and their interaction on species diversity and density of seed banks of grazed sand grasslands (GLMM). Notations: marginally significant effects (*p*<0.1) were denoted with *italics*, significant effects (*p*<0.05) with **boldface**. CWM = community weighted means, LA = leaf area, LDMC = leaf dry matter content, SLA = specific leaf area. Coordinates for C, S and R strategy of a particular species ranges from a theoretical minimum of 0 to a theoretical maximum of 100 (percentage scale). Sum of the three coordinates for a particular species or site are 100. SBT= Social Behaviour Types by Borhidi 1994, *Species of natural and semi natural habitats:* natural pioneers (NP), generalists (G), specialists (S) and natural competitors (C). *Disturbance tolerators* are disturbance tolerators (DT). *Weeds* are the categories weeds (W), adventives (A), ruderal competitors (RC) and adventive competitors (AC). The trends of significantly affected dependent variables are shown in figures and Appendix B. Family-wise error rate was controlled by applying Bonferroni-Holm correction to the raw *p* values within each family of related response variables.

| Characteristics | Livestock type |  | Grazing intensity |  | Livestock type<br>× Grazing<br>intensity |  |
| --- | --- | --- | --- | --- | --- | --- |
| | $F_{1,117}$ | $p_{Holm}$ | $F_{3,117}$ | $p_{Holm}$ | $F_{3,117}$ | $p_{Holm}$ |
| Shannon diversity | 0.748 | 1.000 | 3.266 | 0.143 | *4.038* | *0.054* |
| Evenness | 0.001 | 1.000 | 0.966 | 1.000 | 1.749 | 0.321 |
| *Species richness* |  |  |  |  |  |  |
| Total | 1.780 | 1.000 | 2.691 | 0.247 | **7.208** | **0.001** |
| Short-lived forb | 0.855 | 1.000 | 1.459 | 0.917 | 2.344 | 0.229 |
| Short-lived graminoid | 0.876 | 1.000 | **5.953** | **0.006** | 1.751 | 0.321 |
| Perennial forb | 1.060 | 1.000 | 0.506 | 1.000 | *3.557* | *0.083* |
| Perennial graminoid | 0.889 | 1.000 | 0.013 | 1.000 | *3.534* | *0.083* |
| *Seed bank density* |  |  |  |  |  |  |
| Total | **7.030** | **0.045** | **3.117** | **0.029** | **98.919** | **<0.001** |
| Short-lived forb | 0.015 | 1.000 | **26.154** | **<0.001** | **21.274** | **<0.001** |
| Short-lived graminoid | 0.280 | 1.000 | **6.861** | **<0.001** | **20.347** | **<0.001** |
| Perennial forb | 3.092 | 0.325 | **26.468** | **<0.001** | **62.217** | **<0.001** |
| Perennial graminoid | 0.514 | 1.000 | **11.003** | **<0.001** | **58.055** | **<0.001** |
| *CWM* |  |  |  |  |  |  |
| LA | 2.650 | 0.637 | **8.356** | **<0.001** | **4.485** | **0.010** |
| LDMC | 0.343 | 1.000 | **20.913** | **<0.001** | **5.272** | **0.006** |
| SLA | 1.018 | 1.000 | **12.657** | **<0.001** | **6.404** | **0.002** |
| C coordinate | 0.436 | 1.000 | **5.028** | **0.003** | 1.488 | 0.221 |
| S coordinate | 0.292 | 1.000 | **14.877** | **<0.001** | **5.990** | **0.003** |
| R coordinate | 0.099 | 1.000 | **14.886** | **<0.001** | **8.068** | **<0.001** |
| *SBT* |  |  |  |  |  |  |
| Species of natural and semi natural habitats | **9.285** | **0.009** | **18.418** | **<0.001** | **78.523** | **<0.001** |
| Disturbance tolerators | 1.197 | 0.552 | **13.197** | **<0.001** | **110.046** | **<0.001** |
| Ruderal and weedy species | 0.143 | 0.706 | **85.590** | **<0.001** | **23.171** | **<0.001** |

## 3. Results

### 3.1. Species composition of the vegetation and the seed bank

We identified altogether 192 vascular plant species in the vegetation and the soil seed banks; 137 species were found in the vegetation while 126 species in the SSB (66 species only in the vegetation, 71 species both in the vegetation and the SSB, while 55 species only in the SSB, App. A). In addition, three further species (*Epilobium* sp.*, Oxalis* sp*., Sonchus oleraceus*) emerged, but were considered as contamination because they were found in the control pots and therefore omitted from further analyses. Of the most frequent species, *Anthemis ruthenica*, *Arenaria serpyllifolia*, *Cerastium semidecandrum*, *Cynodon dactylon*, *Erigeron canadensis*, *Myosotis stricta*, *Poa angustifolia*, *Potentilla argentea*, *Potentilla incana*, *Rumex acetosella*, *Scleranthus annuus*, *Veronica arvensis*, *Veronica verna*, and *Vicia lathyroides* were detected both in the vegetation and the seed banks. In the vegetation, *Cerastium pumilum* subsp. *glutinosum*, *Chondrilla juncea*, *Crepis rhoeadifolia*, *Hypochoeris radicata*, and *Spergula pentandra* were not detected in the soil seed bank. In contrast, from the most frequent seed bank species *Eragrostis pilosa*, *Portulaca oleracea*, and hygrophyte species like *Typha angustifolia* and all the detected *Juncus* species were present only in the seed bank (App. A).

We compared the species composition of vegetation and soil seed banks of the grazed grasslands using a PCA ordination (Fig. 3). We found that vegetation and seed bank compositional data were quite well separated from each other. The vegetation of X19-X21 sites formed a relatively distinct point cloud in the ordination and were characterised by a higher frequency of *Jasione montana*. No discrete clustering of vegetation and seed bank composition was detected based on either livestock type or grazing intensity except of the sites X19-X21 which were cattle-grazed ones.

**Figure 3.**
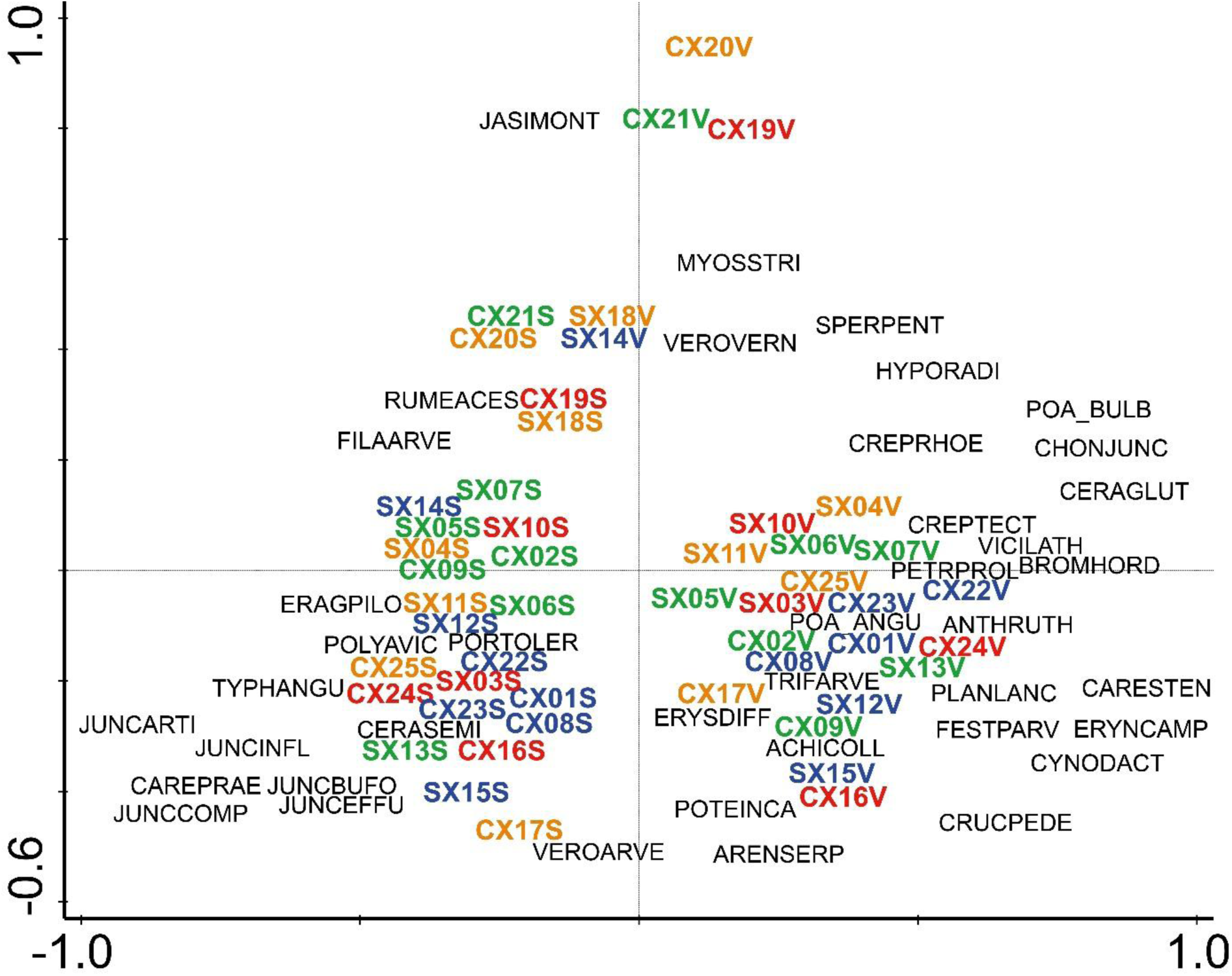
Vegetation and seed banks of the studied 25 sand grassland sites displayed by a PCA ordination. The ordination was based on a presence-absence dataset, where frequency data was used (the number of plots/soil samples per site where the species occurred were summarised, the frequency was 0 when the species was not detected in any plots/soil samples, and 5 when the species was present in all plots/soil samples). The 30 most frequent species either in the vegetation or the seed banks were filtered and added to the ordination diagram (altogether 46 species are shown, 14 frequent species were present both in vegetation and seed banks). The cumulative species variance explained by the four axes of ordination was 46.13. The names of the species follow a four-letter abbreviation of the genus, and a four-letter abbreviation of the species names as follows: ACHICOLL = *Achillea collina*, ANTHRUTH = *Anthemis ruthenica*, ARENSERP = *Arenaria serpyllifolia*, BROMHORD = *Bromus hordeaceus*, CAREHIRT = *Carex hirta*, CAREPRAE = *Carex praecox*, CARESTEN = *Carex stenophylla*, CERAGLUTI = *Cerastium pumilum* subsp. *glutinosum*, CERASEMI = *Cerastium semidecandrum*, CHONJUNC = *Chondrilla juncea*, CREPRHOE = *Crepis rhoeadifolia*, CREPTECT = *Crepis tectorum*, CRUCPEDE = *Cruciata pedemontana*, CYNODACT = *Cynodon dactylon*, ERAGPILO = *Eragrostis pilosa*, ERIGCANA = *Erigeron canadensis*, ERYNCAMP = *Eryngium campestre*, ERYSDIFF = *Erysium diffusum*, FESTPARV = *Festuca valesiaca* subsp. *parviflora*, FILAARVE = *Filago arvensis*, HYPERPERF =*Hypericum perforatum*, HYPORADI = *Hypochoeris radicata*, JASIMONT = *Jasione montana*, JUNCARTI = *Juncus articulatus*, JUNCBUFO = *Juncus bufonius*, JUNCCOMP = *Juncus compressus*, JUNCEFFU = *Juncus effusus*, JUNCINFL = *Juncus inflexus*, MYOSSTRI = *Myosotis stricta*, PETRPROL = *Petrorhagia prolifera*, PLANLANC = *Plantago lanceolata*, POA_ANGU = *Poa angustifolia*, POA_BULB = *Poa bulbosa*, POLYAVIC = *Polygonum aviculare*, PORTOLER = *Portulaca oleracea*, POTEARGE = *Potentilla argentea*, POTEINCA = *Potentilla incana*, RUMEACES = *Rumex acetosella*, SCLEANNU = *Scleranthus annuus*, SPERPENT = *Spergula pentandra*, TRIFARVE = *Trifolium arvense*, TYPHANGU = *Typha angustifolia*, VERBPHLO = *Verbascum phlomoides*, VEROARVE = *Veronica arvense*, VEROVERN = *Veronica verna*, VICILATH = *Vicia lathyroides*. Explanation of the site codes in order of appearance in the abbreviations: first letter- C= cattle or S = sheep then X1-X25 site codes and the last letter is V= vegetation or S= seed banks. Grazing intensities are denoted with different colours: green = 1, blue = 2, orange = 3, and red = 4).

### 3.2. Seed bank density

Altogether 9,507 seeds germinated from the samples; the average SSB density (mean) of sheep-grazed sites was 7,748 seeds/m^2^, while the density was 12,247 seeds/m^2^ in cattle-grazed sites. The 20 most frequent species were the followings (see total seed numbers in the brackets): *Rumex acetosella* (1666), *Filago minima* (592), *Potentilla argentea* (582), *Juncus articulatus* (534), *Carex praecox* (509), *Juncus compressus* (487), *Juncus bufonius* (437), *Cerastium semidecandrum* (419), *Portulaca oleracea* (327), *Veronica verna* (272), *Scleranthus annuus* (258), *Arenaria serpyllifolia* (253), *Jasione montana* (246), *Juncus inflexus* (235), *Conyza canadensis* (214), *Juncus effusus* (179), *Potentilla arenaria* (134), *Anthemis ruthenica* (124), *Ambrosia artemisiifolia* (120), *Verbascum phlomoides* (110). See further data on the seed densities in App. A.

### 3.3. Effect of livestock type on the seed bank

Livestock type had a significant main effect on the total SSB density and the SSB density of species of natural and semi-natural habitats (Tab. 2). The main effect of livestock type was not significant on any of the 15 most abundant species in the SSB (Tab. 3., App. B, lower scores for sheep-grazed sites regardless to grazing intensity). Regarding total SSB density, the main effect of livestock type was significant; and we found higher seed densities in cattle-grazed sites with a striking difference at the second and particularly at the fourth level of grazing intensity (Fig. 4A). The SSB density of species of natural and semi-natural habitats was significantly affected by the main effect of livestock type, with typically higher scores for cattle-grazed sites, especially at the fourth grazing intensity level (Tab. 2, Fig. 5A). The main effect of livestock type was not significant on the CWM of CSR coordinates (Tab. 2). Sites grazed by cattle or sheep were well separated in the CSR triangles only at the first intensity level (App. C).

**Figure 4.**
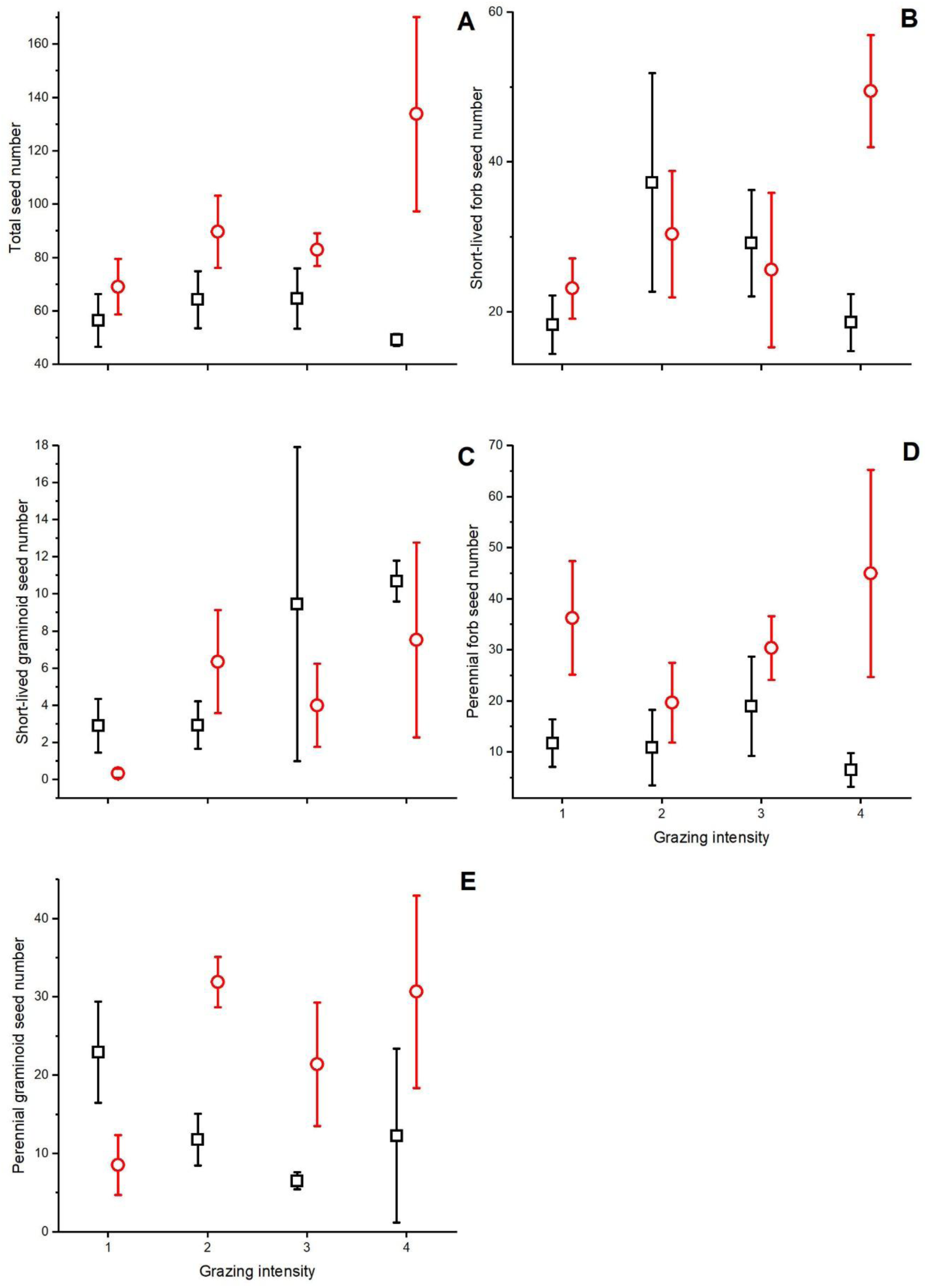
Effect of livestock type and grazing intensity on total seed bank density (A), and on the seed bank density of morpho-functional groups: short-lived forbs (B), short-lived graminoids (C), perennial forbs (D), and perennial graminoids (E). Total seed bank density of sheep-grazed (black square) and cattle-grazed (red circle) grasslands (mean ± SE). For grazing intensity levels see Table 1. The number of seeds per plot (6 soil cores) is shown in the figure, one detected seed corresponds to 133 seeds/m^2^ seed density in the upper ten centimetres of soil.

**Figure 5.**
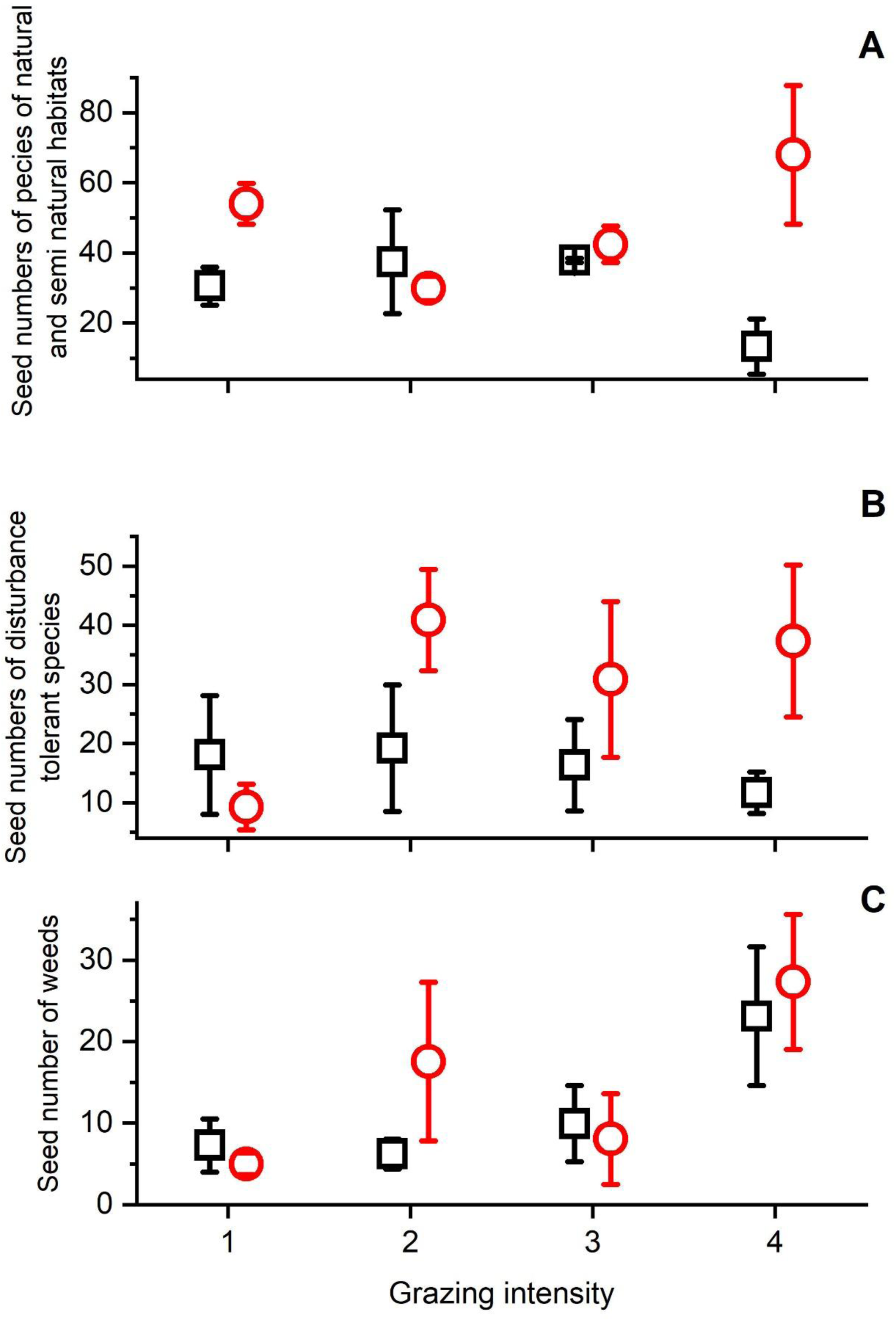
Effect of grazing intensity, livestock type and their interaction on density of seed banks of Social Behaviour Types (SBT) groups of grazed sand grasslands (GLMM, mean ± SE). The studied groups were: i) *Species of natural and semi-natural habitats:* natural pioneers (NP), generalists (G), specialists (S) and natural competitors (C). ii) *Disturbance tolerators* (DT). iii) *Weeds* are the categories weeds (W), adventives (A), ruderal competitors (RC) and adventive competitors (AC) (Borhidi, 1995). The test results can be found in Table 2.

**Table 3.** Effect of grazing intensity, livestock type and their interaction on the density of soil seed banks of species with the highest cumulative seed numbers on the studied grazed sand grasslands (GLMM; species with >200 viable seeds detected in the 25 sites in total were tested). Notations: marginally significant effects (p<0.1) were denoted with *italics*, significant effects (*p*<0.05) with **boldface**. Family-wise error rate was controlled by applying Bonferroni-Holm correction to the raw *p* values within each family of related response variables.

|  | Livestock type |  | Grazing intensity |  | Livestock type ×<br>Grazing intensity |  |
| --- | --- | --- | --- | --- | --- | --- |
| Seed bank density | $F_{1,117}$ | $p_{Holm}$ | $F_{3,117}$ | $p_{Holm}$ | $F_{3,117}$ | $p_{Holm}$ |
| *Rumex acetosella* | 6.232 | 0.209 | **24.431** | **<0.001** | **50.362** | **<0.001** |
| *Filago minima* | 3.748 | 0.774 | 1.151 | 1.000 | 1.620 | 0.754 |
| *Potentilla argentea* | 0.183 | 1.000 | **6.708** | **<0.001** | **38.585** | **<0.001** |
| *Juncus articulatus* | 0.202 | 1.000 | **10.700** | **<0.001** | **26.160** | **<0.001** |
| *Carex praecox* | 0.185 | 1.000 | 2.985 | 0.205 | **6.623** | **0.003** |
| *Juncus compressus* | 3.176 | 1.000 | 0.163 | 1.000 | **8.336** | **<0.001** |
| *Juncus bufonius* | 0.025 | 1.000 | 2.033 | 0.565 | **18.805** | **<0.001** |
| *Cerastium semidecandrum* | 0.015 | 1.000 | 0.502 | 1.000 | 3.623 | 0.091 |
| *Portulaca oleracea* | 0.015 | 1.000 | **35.027** | **<0.001** | 3.147 | 0.139 |
| *Veronica verna* | 1.004 | 1.000 | 3.322 | 0.156 | 0.749 | 1.000 |
| *Scleranthus annuus* | 0.080 | 1.000 | **25.767** | **<0.001** | **8.571** | **<0.001** |
| *Arenaria serpyllifolia* | 2.143 | 1.000 | **10.324** | **<0.001** | 1.492 | 0.754 |
| *Jasione montana* | 1.472 | 1.000 | 0.107 | 1.000 | 0.104 | 1.000 |
| *Juncus inflexus* | 1.390 | 1.000 | **4.847** | **0.028** | **5.351** | **0.012** |
| *Erigeron canadensis* | 0.036 | 1.000 | **4.871** | **0.028** | **20.905** | **<0.001** |

### 3.4. Effect of grazing intensity on the soil seed bank

Grazing intensity had a significant main effect on many of the studied variables except for most diversity measures and the species richness characteristics of morpho-functional groups (Tab. 2). Total species richness was higher at the second level of grazing intensity compared to the first level (Fig. 6A). While Shannon diversity was higher at the second grazing intensity level compared to the first one (Fig. 6B), no clear pattern was found for evenness (Fig. 6C). Species richness of short-lived graminoids was significantly lower at the first level compared to the other levels, but the richness of other functional groups were not affected by the main effect of grazing intensity (Tab. 2, Fig. 7). The seed density of all morpho-functional groups was significantly affected by the main effect of grazing intensity (Tab. 2, Fig. 4). The seed density of short-lived forbs (Fig. 4B), perennial forbs (Fig. 4D), and perennial graminoids (Fig. 4E) was much higher in cattle-grazed sites at the fourth level of grazing intensity compared to the other levels. LA was significantly higher at the fourth level of grazing intensity compared to the other levels (Fig. 8A). Between the second and fourth levels of grazing intensity, LDMC decreased significantly (Fig. 8B), while SLA significantly increased along the levels of grazing intensity (Fig. 8C). Grazing intensity significantly affected the CWMs of both C, S, and R coordinates. The CWM of C coordinates was significantly higher at the fourth level of grazing intensity compared to the other levels (Fig. 8D). In the case of the CWM of R and S coordinates, the second level of grazing intensity was significantly different compared to the other levels: at the second intensity level, the CWM of S coordinates was significantly higher, while that of R coordinates was significantly lower (Fig. 8E and 8F). Grazing intensity also significantly affected the seed density of all three categories of Social Behaviour Types (Tab. 2). The largest difference in mean seed bank abundance between sheep-grazed and cattle-grazed sites occurred at the second grazing intensity level for ruderal and weedy species, while for natural and semi-natural species the greatest difference was observed at the fourth grazing intensity level (both cases higher figures for cattle-grazed sites, App. B). For the disturbance-tolerant species, higher values for cattle-grazed sites were typical at the second, third and fourth intensity levels. Considering the 15 most abundant species, the SSB densities of eight species were affected by the main effect of grazing intensity (Tab. 3). The responses of species were highly species-specific and strong interactions were detected (App. B).

**Figure 6.**
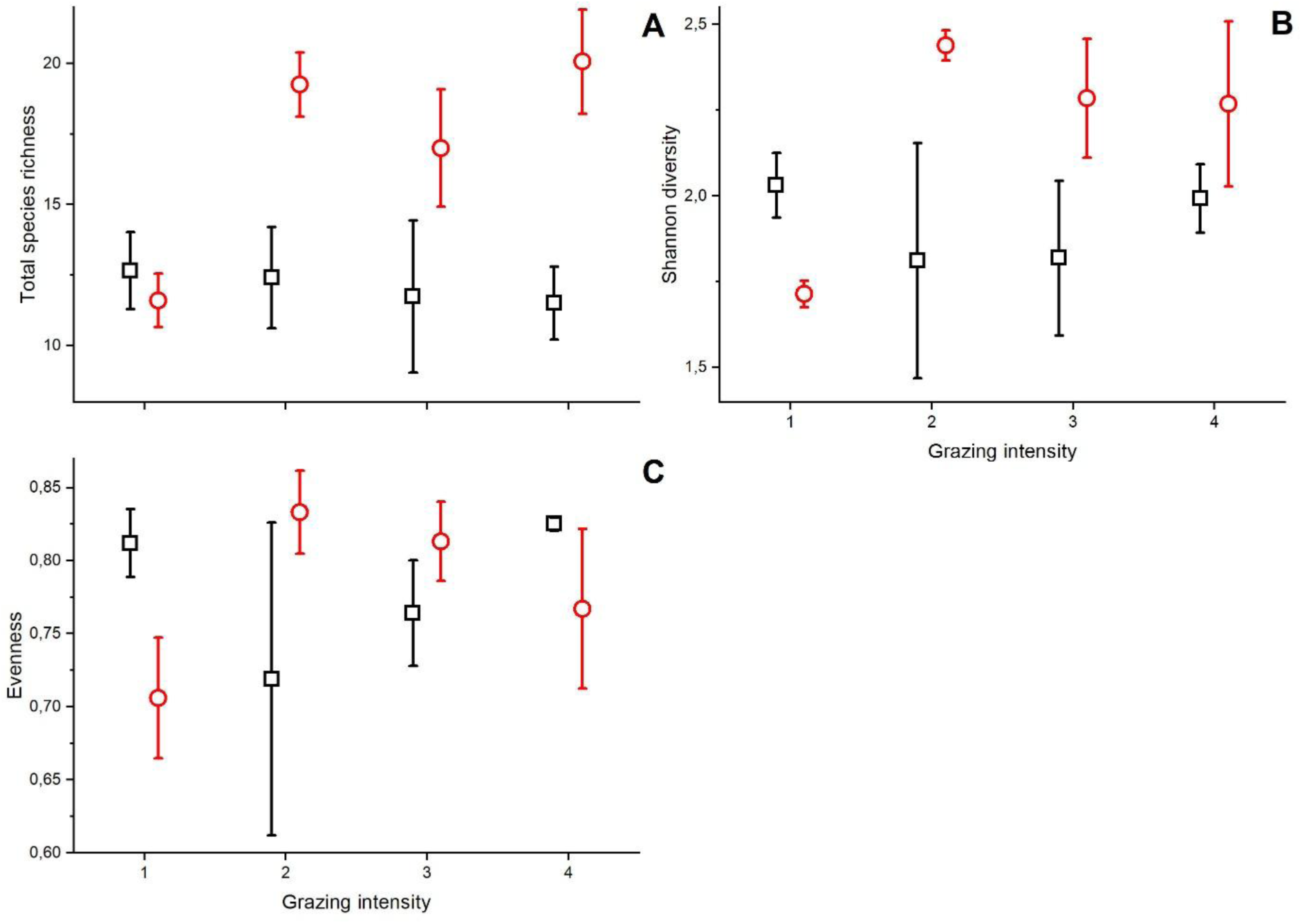
Total species richness (A), Shannon diversity (B) and Evenness (C) along the increasing grazing intensity (mean ± SE) in sheep-grazed versus cattle-grazed sites. Significant differences (GLMM and Tukey test, p<0.05, see detailed statistics in Table 1) are indicated by different superscripted letters. For the explanation of grazing intensity levels please see Table 1.

**Figure 7.**
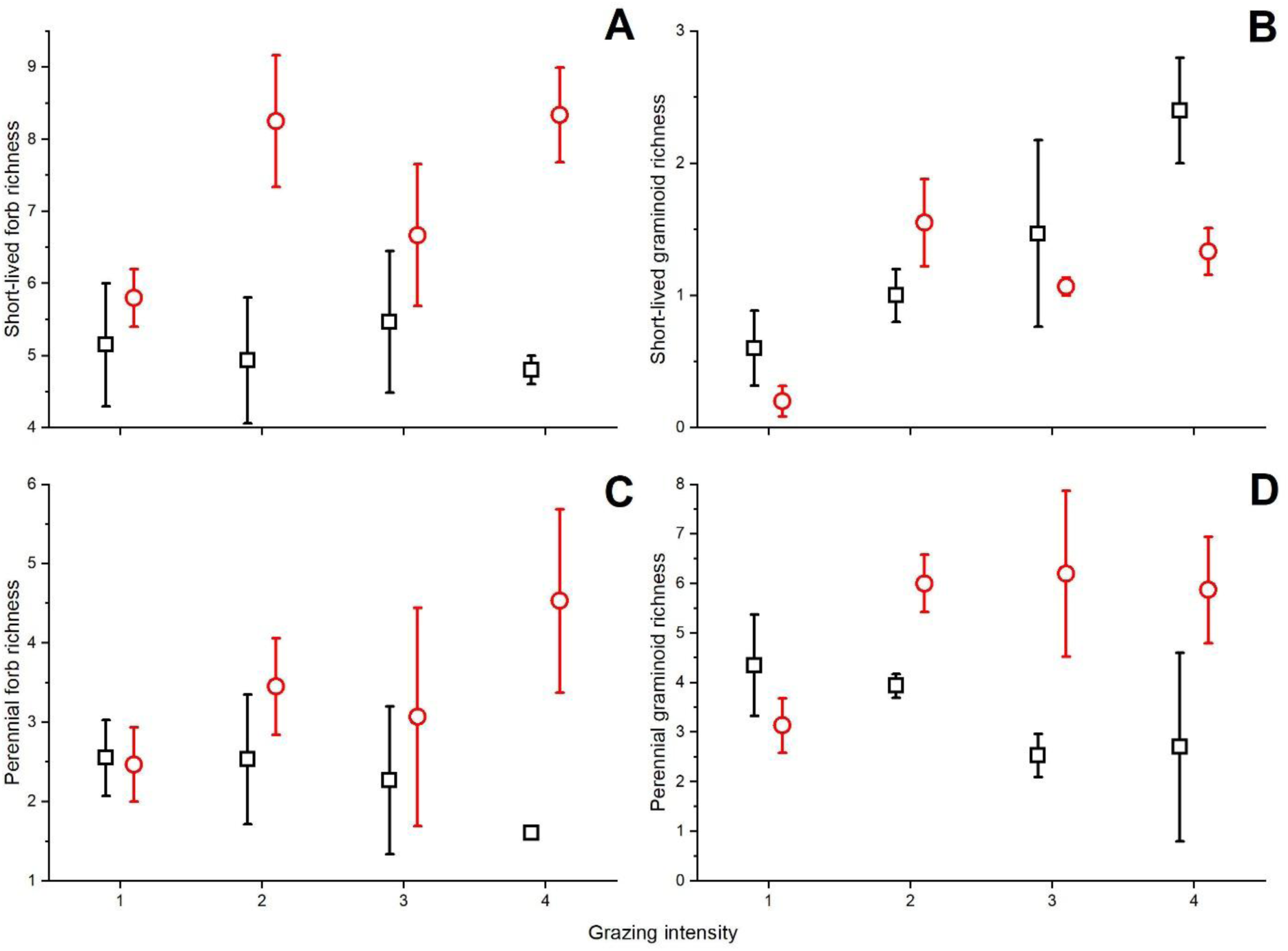
Species richness of short-lived graminoids (A), short-lived forbs (B), perennial graminoids (C) and perennial forbs (D) along the increasing grazing intensity (species per plot mean ± SE) in sheep-grazed versus cattle-grazed sites. Significant differences (GLMM and Tukey test, *p*<0.05, detailed statistics see in Table 1) are indicated by different superscripted letters. For the explanation of grazing intensity levels please see Table 1.

**Figure 8.**
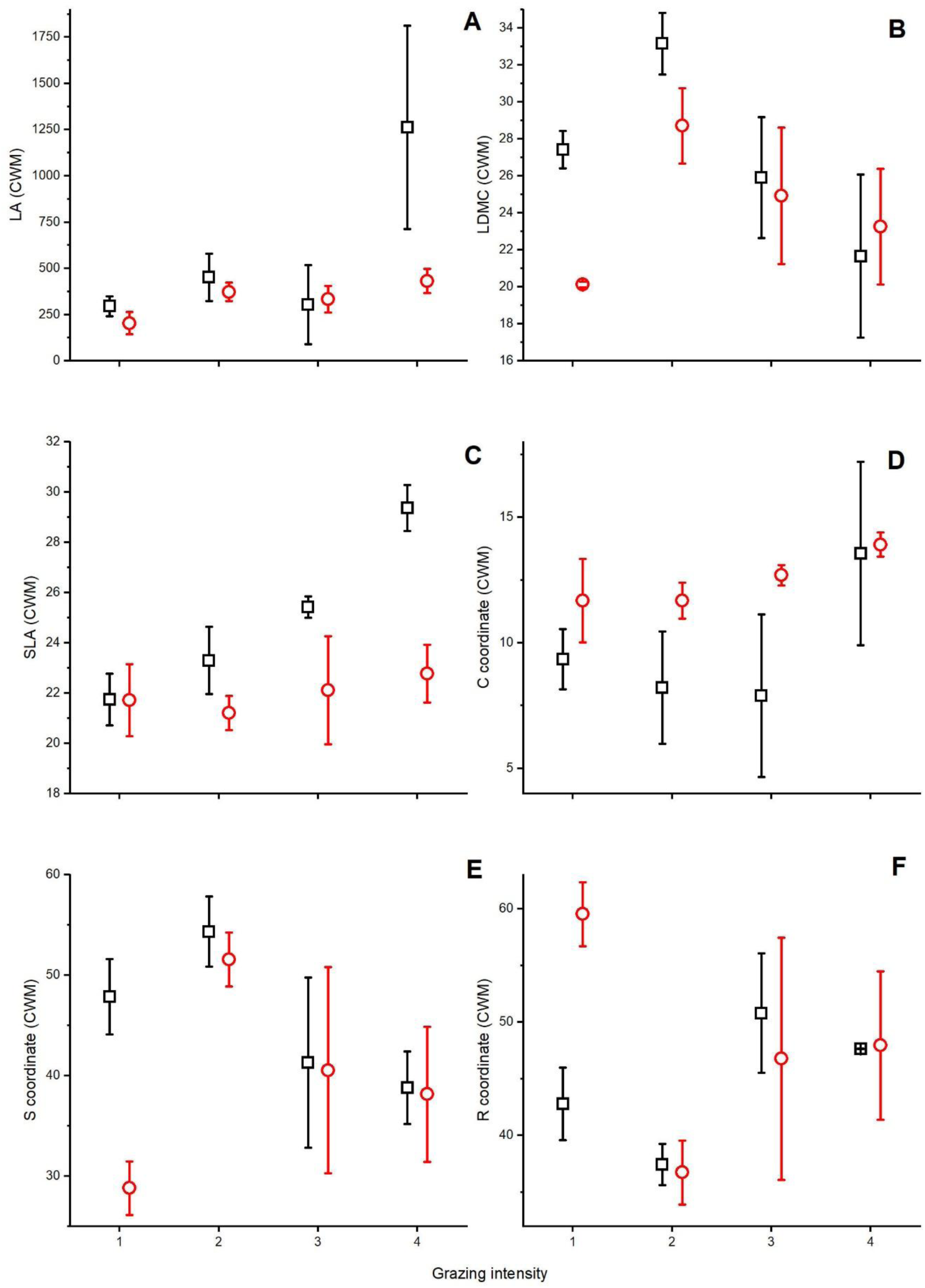
Community-weighted means of leaf area (LA; A), leaf dry matter content (LDMC; B), and specific leaf area (SLA; C), and CSR coordinates: C (D), S (E), and R (F) along the increasing grazing intensity (mean ± SE) in sheep-grazed versus cattle-grazed sites. Significant differences (GLMM and Tukey test, *p*<0.05, detailed statistics see in Table 1) are indicated by different superscripted letters. For the explanation of grazing intensity levels please see Table 1.

### 3.5. Effect of interaction of grazing intensity and livestock type on the soil seed bank

We found that most cases where the main effect grazing intensity on dependent variables was significant, also the interaction effect of livestock type and grazing intensity was found to be significant (Tab. 2). The total species richness was significantly affected by the interaction of livestock type and grazing intensity, displaying an increasing trend with increasing grazing intensity at cattle-grazed sites and a decreasing trend for sheep-grazed sites (the total richness of the seed bank was markedly higher for cattle-grazed sites than for sheep-grazed ones at grazing intensity levels 2–4). The interaction of livestock type and grazing intensity also had a significant effect on the seed bank density of all morpho-functional groups and on all leaf traits and SBT groups (Tab. 2). Increasing SSB density values were detected along an increase in grazing intensity in cattle-grazed sites, while mostly decreasing or unaltered values were found with increasing levels of sheep grazing (Fig. 4). The largest differences were detected for LA and SLA, with increasing values with increasing grazing intensity at sheep-grazed sites, but with unchanged values along the intensity gradient in cattle-grazed ones (Fig. 8). The interaction of livestock type and grazing intensity had a significant effect on the CWMs of S and R coordinates (Tab. 2), but the changes for S and R coordinates for the different livestock types along the intensity gradient were not that evident as in the case of some other variables mentioned above. From the 15 most abundant species, the seed bank density of 9 species was affected by the interaction of intensity and livestock type (Tab. 3). The SSB density of most *Juncus* species increased with increasing grazing intensity in case of cattle grazing (App. B). The seed bank density of some abundant small-sized short-lived species like *Cerastium semidecandrum* or *Scleranthus annuus* showed different patterns for cattle and sheep grazing. While the abundance of *C. semidecandrum* increased in cattle-grazed sites and decreased in sheep-grazed ones with increasing grazing intensity (marginally significant interaction effect), that of *S. annuus* showed an opposite trend peaking at the most intensively grazed sheep pastures (Tab. 3, App. B).

## 4. Discussion

### 4.1. Effect of livestock type on the SSB

We partly confirmed our first hypothesis that sheep-grazed sites have lower SSB diversity and density compared to cattle-grazed ones. The total density of SSBs (at all intensity levels) and the density of natural and semi-natural species (at all but one intensity level) were lower in sheep-grazed sites compared to cattle-grazed ones. However, the diversity metrics and the SSB density of functional groups and individual species were not affected by the main predictor ‘livestock type’. However, the interaction of grazing intensity and livestock type was significant also for the total species richness of the soil seed banks, and on the seed bank density of all the studied morpho-functional groups and functional species groups (Tab. 2). This means that the values of these dependent variables showed a different pattern to the increase of grazing intensity in cattle and sheep-grazed sites. Considering the species-specific results, it was also found that from the 15 most abundant species, none of the species was affected by the main effect of livestock type. According to Shi et al. (2022), most studies addressing grazing and seed bank have no information about the livestock type. Studies addressing grazing and vegetation provide more information in this respect. Metera et al. (2010) and Jerrentrup et al. (2015) found that sheep have a higher selectivity for forbs, which leads to higher functional and taxonomical diversity in the vegetation for cattle grazing (Tóth et al., 2016). We could not confirm the overall higher diversity for SSB in our study, although the species richness of cattle-grazed sites exceeded that of sheep-grazed ones from the 2^nd^ to the 4^th^ level of grazing intensity (Fig. 6). However, total seed density was significantly higher in cattle-grazed sites, which is probably due to cattle being less selective for forbs. Moreover, sheep consume small individual plants and bite very close to the ground using their incisors in contrast to cattle (Metera et al., 2010; Jerrentrup et al., 2015), which presumably opens more gaps in the vegetation, and increased germination in the created gaps can reduce the SSB density. Cattle are less selective for individual plants but selective for patches with higher biomass due to their feeding strategy as well: cattle use their tongue wrapping the plants (Rook et al., 2004; Jerrentrup et al., 2015). Lower selectivity of cattle can result in more plants ripening seeds, for example, short-lived forbs ripening small-sized seeds in large amounts (Coomes & Grubb, 2003). We found that the pattern in the seed density of short-lived forbs closely resembles the total seed density pattern along grazing intensity levels in case of cattle grazing. It suggests that both the seed density of short-lived forbs and total seed density are positively affected by the proximity of frequently visited places (the second and the fourth levels of grazing intensity). Despite the patterns found, seed density of short-lived forbs was not significantly affected by livestock type only in interaction with grazing intensity. However, it does not mean that cattle grazing compared to sheep grazing increases the seed density of short-lived forbs in general, because the effect was highly species-specific and some short-lived species (e.g., *Scleranthus annuus*) benefitted from the increasing intensity of sheep grazing. Sheep grazing was found to increase the biomass of short-lived, weedy forbs in sand grasslands when grazing intensity was higher, especially when frequently visited places were closer (Kovacsics-Vári et al., 2024). In addition, grazing-driven changes in biomass can increase the similarity between aboveground vegetation and SSB in the long run (Sanou et al., 2018; Zida et al., 2020). This is probably because the formerly mentioned feeding strategy makes cattle less selective, not only for forbs (Rook et al., 2004; Jerrentrup et al., 2015). Our findings draw the attention to the careful planning of grazing management (grazing intensity, livestock type, and probably the arrangement of grazing lands) if we wish to prevent the accumulation of seeds of species which are undesirable from a conservation point of view.

### 4.2. Effect of grazing intensity on the SSB

We found that grazing intensity had a significant main effect on some of the diversity measures, on the SSB density of all plant functional groups, on the leaf traits, on the CSR strategy composition and on the species groups formed using SBT as well. On the other hand, only the total seed bank density and the density of species of natural and semi-natural habitats were affected by the main effect of ‘livestock type’. Thus, we confirmed the second hypothesis that the species composition, diversity, and density of the SSBs are more strongly affected by grazing intensity than by livestock type. To our knowledge, no previous study compared the effect of livestock type and grazing intensity on the SSB. Former studies addressed the aboveground vegetation and had contrasting results; some previous studies conducted in sand grasslands agree with our results (Kovacsics-Vári et al., 2023) while some others conducted in different grassland types contradict them (Tóth et al., 2016 – alkaline grasslands; Rodriguez et al., 2023 - mountain grasslands). However, it is important to stress that in most cases a significant effect of the interaction of livestock type and grazing intensity was also detected. Considering the above-mentioned results, we clearly demonstrated that livestock type and grazing intensity are both important drivers of the diversity of vegetation and seed banks. It is also important to mention that several former studies have demonstrated that effects of grazing intensity and livestock type are also highly dependent on the type of the subjected grassland (Rook et al., 2004; Fraser et al., 2022; Jiang & Wang, 2022; Török et al., 2024).

Regardless of livestock type, former studies found that the amount of green biomass and litter was lower at higher grazing intensities (Magnano, et al., 2019; Kovacsics-Vári et al., 2023), and dominant species were supressed by grazers (Salguero-Gómez, 2017) which promotes the seed bank formation of morpho-functional groups consisting of subordinated species. This explanation can be confirmed by our study as species richness and Shannon diversity were higher at the second level of intensity compared to the first level. According to the meta-analysis by Shi et al. (2022), light grazing can increase species richness in the SSB. We found that both species richness and Shannon diversity were significantly higher at the second level of grazing intensity than at the first level and we detected no further increase in these variables at higher intensity levels that can confirm Shi et al. (2022).

In contrast, we cannot confirm the result by Shi et al. (2022) that SSB density is higher at lower intensity of grazing. When grazing intensity was higher, SSB density was found to be lower both in mesic and dry habitats including not only grasslands (Shi et al, 2022). These findings were confirmed (Zhao et al., 2011) as the consumption of plants can reduce the number of seeds. In contrast, seed production can also be higher when grazing intensity is higher (Grime, 2002; Klimkowska et al., 2009). In our study, grazing intensity was found to be a stronger driver of the SSB density than livestock type, but this may only be the case in the selected type of acidic sand grasslands. When grazing intensity is lower, there is no intensive consumption of plants, hence seeds are saved (Zhao et al., 2011) but the amount of accumulated litter is also higher (Magnano, et al., 2019; Kovacsics-Vári et al., 2023). As litter can act as a seed trap (Ruprecht & Szabó, 2012), litter accumulation at low grazing intensity may hinder seeds from being incorporated into the soil and therefore entering the SSB.

The assessment of grazing intensity can differ from study to study, which affects the interpretation of the results (Rodriguez et al., 2023). For example, we considered the LU/ha and also the proximity to frequently visited places (a combination of intensity expressed in LU/ha and the frequency of visits). In our study, when the distance to frequently visited places is lower (second and fourth levels of grazing intensity), the seed bank density of short-lived forbs is higher. In contrast, the density of perennial forbs is higher when the distance to frequently visited places is higher (first and third levels of grazing intensity). Therefore, we found fluctuations for perennial and short-lived forbs along the grazing intensity levels. According to the meta-analysis of Díaz et al. (2007), the abundance of short-lived forbs increases in the aboveground vegetation when grazing intensity is higher, while that of perennial forbs decreases. A meta-analysis of SSB found that short-lived forbs were not affected by grazing, while the density of perennial forbs was decreased by grazing (Shi et al., 2022). In contrast, Li et al. (2024) found that perennial forbs were not affected while the density of short-lived forbs was higher at higher grazing intensity levels (at a soil depth of 0–5 cm). The higher density of short-lived forbs was also confirmed by Dreber & Esler (2011). In our study region (Nyírség, Hungary), the aboveground biomass of sand grasslands was also studied along a gradient of grazing intensity, but contrary to our results, no contrasting effects of grazing on short-lived and perennial forbs were found (Kovacsics-Vári et al., 2024). This confirms the different role of certain species or functional groups in seed bank and in vegetation (Klimkowska et al., 2009).

Surprisingly, perennial graminoids were represented with the highest seed bank density at the highest level of grazing intensity in cattle-grazed sites (Fig. 4E). The opposite was found in studies addressing either aboveground vegetation or seed bank (see e.g., Chu et al., 2019; Li et al., 2024). This contradiction can be explained by the perennial *Juncus* spp. which represent a high seed density in our studied sites (see Appendix A). In mesic to wet grasslands in years with high rainfall and/or after a high decrease in vegetation cover (e.g. by burning), *Juncus* species can emerge from the soil seed banks (where they are likely abundant) with a higher success in the gaps formed and produce a high number of persistent seeds. *Juncus* species can be found with high densities in wet grasslands (Bossuyt & Honnay, 2008). In the studied types of grasslands, it is more likely that seeds of *Juncus* species, together with other small-seeded hygrophytes are dispersed by wind or by grazing animals from deep-lying dune slacks covered by hygrophyte vegetation (Matus et al., 2005).

### 4.3. Effect of grazing intensity and livestock type on leaf traits and CSR strategies

We partially confirmed the third hypothesis that the leaf trait and CSR strategy composition of the SSB are highly affected both by livestock type and grazing intensity. All variables were significantly affected by the main effect of grazing intensity, while no variable was affected by the main effect of livestock type. However, livestock type was visually well separated in the CSR triangle at the first grazing intensity level. According to Pierce et al. (2013), LA determines the C coordinates, LDMC the S coordinates, and SLA the R coordinates, but comparing the CWMs of SLA and R coordinates, the trends are not similar. SLA itself can well reflect the level of grazing intensity in our study. In contrast, SLA has different responses to disturbance severity (LU/ha) and frequency (proximity to frequently visited places) as it was found by Herben et al. (2018). Although R coordinates do not show the same patterns as SLA does, but R coordinates had the highest values in the CSR triangles. Therefore, these results might suggest that CSR strategies indicated grazing disturbance in the SSB quite well. However, the changes of S and R coordinates with grazing intensity are not straightforward. For example, species belonging to the competitive strategy type have large LA and are good competitors when resource availability is stable (Pierce et al., 2013), therefore, we would expect a higher CWM of C coordinates at lower levels of grazing intensity, since circumstances are more beneficial under light disturbances (Westoby et al., 1999; He et al., 2021). Furthermore, species belonging to the ruderal strategy type have a good regeneration ability, thus, they regrow and reproduce soon after the disturbance (Westoby et al., 1999; Pierce et al., 2013). Therefore, we would expect the CWM of R coordinates to be higher at higher levels of grazing intensity. The reason behind the lack of a clear trend may be the variability of plant species at community level because plants have various responses to disturbance and stress. For example, (i) large-leafed, unpalatable species could increase the CWM of C coordinates at higher grazing intensity. (ii) Considering stress, shade- and heat-tolerant species may be favoured by different levels of grazing intensity. The latter idea supports McIntosh-Buday et al. (2024) who found that CSR strategies are too robust for comparing stress in saline and loess vegetation. To better interpret the response of CSR strategies to grazing management, a thorough knowledge of species is needed (e.g., how percentage values of strategies of species relate to frequently visited places/grazing frequency and stocking rate). In addition, further traits and functional groups should be included into the trait-based analyses.

### 4.4. The effect of the interaction of livestock type and grazing intensity on the SSB

The most interesting results were the interactions between livestock type and grazing intensity. Surprisingly, many studied dependent variables were significantly affected by the interaction of grazing intensity and livestock type. The difference in total species richness and seed bank density and in the richness and density of perennial species in the seed banks between cattle and sheep-grazed sites became more pronounced at the higher grazing intensity levels on the benefit of cattle-grazed sites. This trend was also visible for the species of natural and semi-natural habitats and for disturbance tolerant species (Appendix B). These findings are in line with results found for aboveground vegetation comparing sheep and cattle grazing in other types of grasslands. For example, Tóth et al. (2016) found lower richness and diversity in sheep-grazed pastures compared to cattle grazing in short-grass steppes. The lower taxonomic richness and lower amount of forb species were also found by other authors for sheep-grazed pastures (Pykälä, 2005; Jerrentrup et al., 2015). The detected differences between cattle and sheep grazing are explained by the higher selectivity of sheep for subordinated forbs in grassland communities (Dumont et al., 2011). However, for example Kovacsics-Vári et al. (2023) found no effect of livestock type on the species richness and diversity of sand grassland vegetation, but the interactions between livestock type and grazing intensity were found to be also significant. These discrepancies can be explained by the species-specific effects of grazing, found also in our study that species in the soil seed bank displayed different responses to the interaction of livestock type and grazing. While for many abundant species, we found higher SSB densities in cattle-grazed pastures or detected an increasing trend of their seed bank densities along an increasing intensity of cattle grazing (i.e. *Rumex acetosella*, *Juncus* species, or *Cerastium semidecandrum*), some other species displayed the opposite trend (*Scleranthus annuus*). This highlights the fact that grazing responses should be carefully considered and compared also at the species level.

## 5. Conclusions and implications for conservation

Importantly, we found that seed bank characteristics were more strongly affected by the main effects of grazing intensity than by live-stock type, but i) seed density was rather livestock-type dependent, especially closer to the frequently visited places and ii) the interaction between grazing intensity and livestock type significantly affected many of the studied variables. Although the seed density of short-lived forbs was not significantly affected by the main effect of livestock type, their seed density was significantly higher in sites that are closer to frequently visited places, especially in case of cattle grazing at the highest level of grazing intensity. These findings raise the question of the adequate grazing frequency and the good arrangement of the drinking points, resting points and barns in grazing lands, especially ones that are small and/or situated near human-made habitats facilitating the encroachment of weeds and invasive species. We have also found that sheep grazing tended to decrease the total seed bank density especially at higher grazing intensities compared to cattle grazing, and this effect was also valid for the seed bank density of species of natural and semi-natural habitats. Also, with increasing grazing intensity of sheep, the total species richness and the Shannon diversity showed a decreasing trend compared to cattle grazing. Thus, high intensity sheep grazing should be avoided in sites rich in forbs. We emphasise that for a sustainable grazing management and to achieve nature conservation goals, grazing intensity and livestock type need to be considered simultaneously.

## Acknowledgements

We are thankful to Demeter, L., Kovács, Z., Patalenszki, N., Pompola, K., Széll, L. (rangers of Hortobágy National Park) for providing information on grazing. Takács, A. (University of Debrecen) supported us with species identification of transplants. We thank the Botanical Garden of the University of Debrecen for providing space for the seedling emergence. We thank to Odame, Amo, D., Godana, Duba, S., Boros, B., Detvai, P., Espinoza, Ami, F.D., Gutiérrez, Perez, D.I., Fekete, T.B., Kasumov, B., Kovács, G., Kumagai, Y., Njoroge, B.W., Papp, SZ., San Martín, Rojas, G.A., Takaró, P., (University of Debrecen) for the essential background work. Kovacsics-Vári, G., McIntosh-Buday, A., and Törő-Szijgyártó, V. were supported by the ÚNKP-23-3-I/II. New National Excellence Program of the Ministry for Culture and Innovation from the source of the National Research, Development and Innovation Found. Sonkoly J. was supported by Hungarian Academy of Sciences, Bolyai János Scholarship (BO/00587/23/8). The research was supported by the National Research, Development and Innovation Office (K137573, PD137747, and KKP144068). This project has received funding from the HUN-REN Hungarian Research Network. The authors were also supported by the University of Debrecen Program for Scientific Publication during manuscript writing. The authors are thankful for the comments and corrections provided by Jürgen Dengler and two anonymous reviewers to the earlier draft of the paper.

## CRediT authorship contribution statement

**Gergely Kovacsics-Vári**: Conceptualization, Data curation, Formal analysis, Investigation, Visualization, Writing – original draft, Writing – review and editing; **Judit Sonkoly**: Conceptualization, Funding acquisition, Investigation, Writing – original draft, Writing – review & editing; **Katalin Tóth**: Data curation, Investigation, Supervision, Validation, Writing – review and editing; **Andrea McIntosh-Buday**: Data curation, Investigation, Project administration; **Luis Roberto Guallichico Suntaxi**: Data curation, Investigation; **Szilvia Madar**: Data curation, Investigation; **Patricia Elizabeth Díaz Cando**: Data curation, Investigation, Writing – review and editing; **Viktória Törő-Szijgyártó**: Data curation, Investigation, Writing – review and editing; **Béla Tóthmérész**: Conceptualization, Investigation, Writing – review and editing; **Péter Török**: Conceptualization, Data curation, Formal analysis, Funding acquisition, Investigation, Methodology, Project administration, Resources, Supervision, Visualization, Writing – original draft, Writing – review and editing

## Declaration of Generative AI and AI-assisted technologies in the writing process

No generative AI or AI-assisted technologies were used in the research process or writing of this manuscript.

## Declaration of competing interest statement

The authors declare that they have no known competing financial interests or personal relationships that could have appeared to influence the work reported in this paper.

## Data availability

The authors in case of acceptance, will make available all underlying data in an open access repository.

## Appendices

**Appendix A.** Vegetation and seed banks of the studied sand grasslands sites (one germinated seedling in the appendix table correspondence with approximately 27 seeds/m^2^ density, pooled data of five plots, altogether with 30 soil cores).

**Appendix B.** Responses of the seed bank density of the most abundant species to grazing intensity, livestock type and their interaction (GLMM, mean ± SE).

**Appendix C.** The CSR strategy composition of the seed banks of sheep-grazed (black squares) and cattle-grazed (red circles) grasslands.

## Notes

### Competing Interest Statement

The authors have declared no competing interest.

