## Appendix B for "High-intensity sheep grazing impoverishes soil seed banks in sand grasslands"

**Appendix B.** Responses of the seed bank density of most abundant species to grazing intensity, livestock type and their interaction (GLMM, mean ± SE). Statistical details see in Table 3. Cattle grazed sites are marked with red circles, sheep with black squares. Number of seeds per plot (6 soil cores) is shown in the figure, one detected seed corresponds with 133 seeds/m$^2$ seed density in the upper ten centimetres of soil.

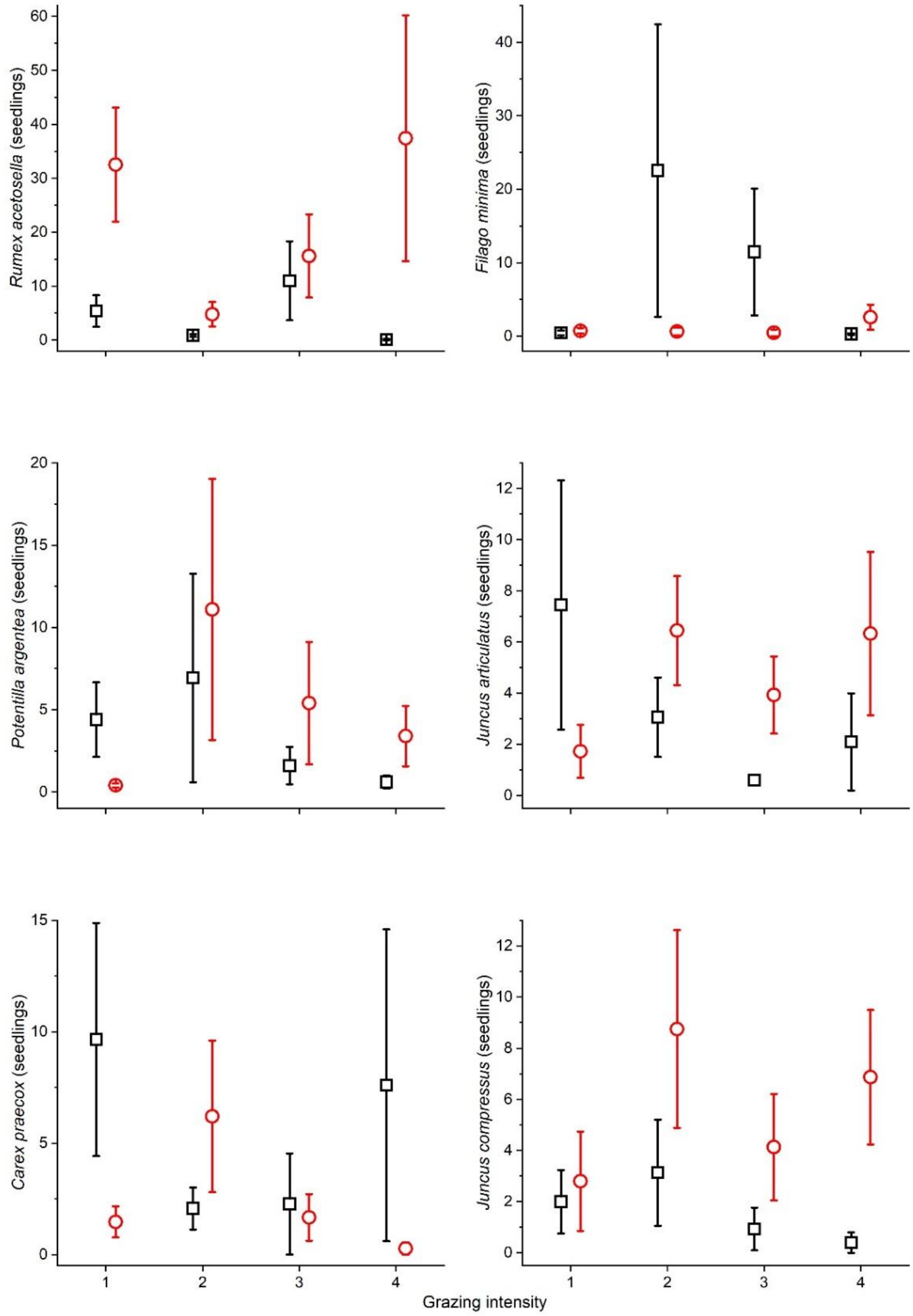

**Appendix B.** Continued.

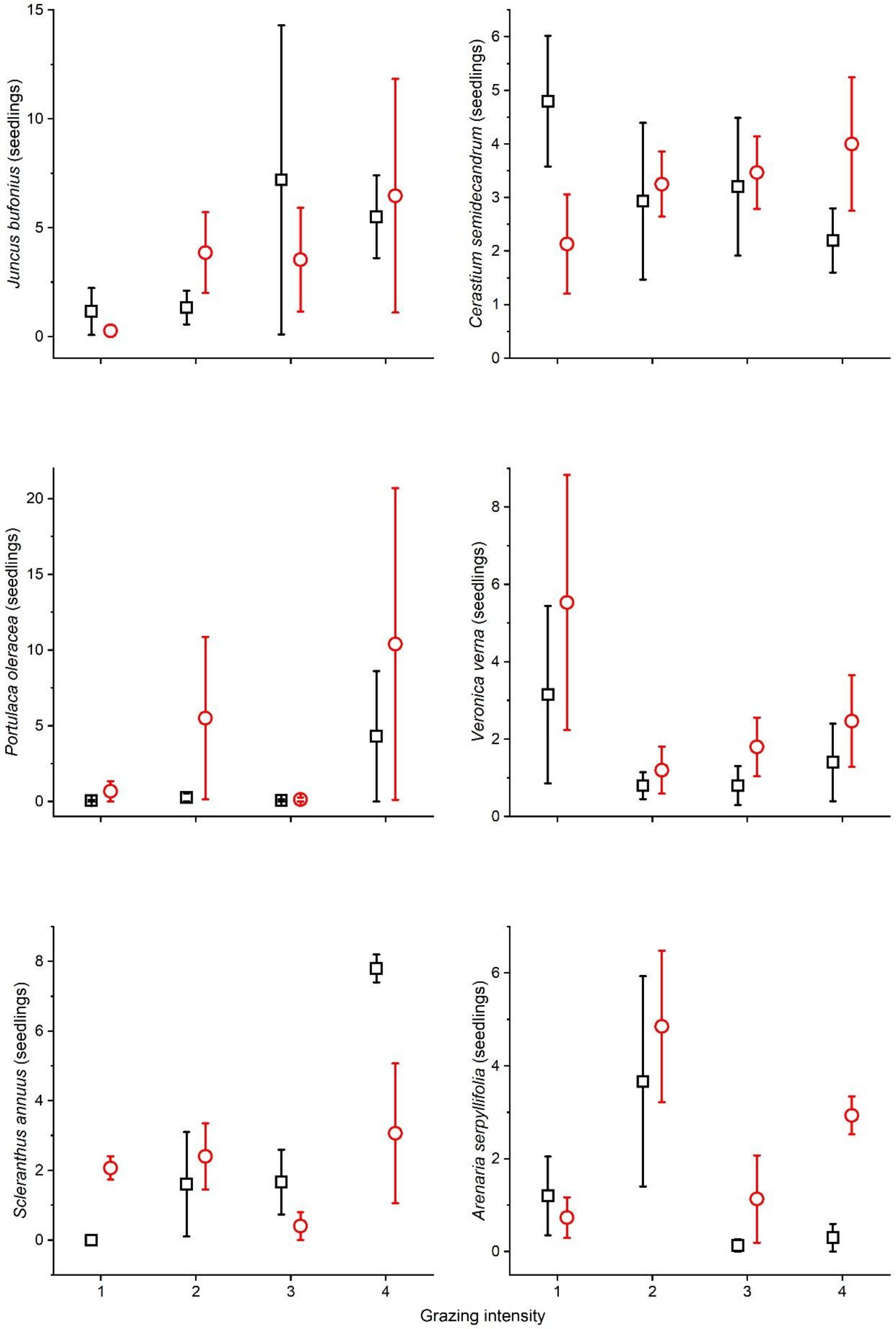

**Appendix B** Continued.

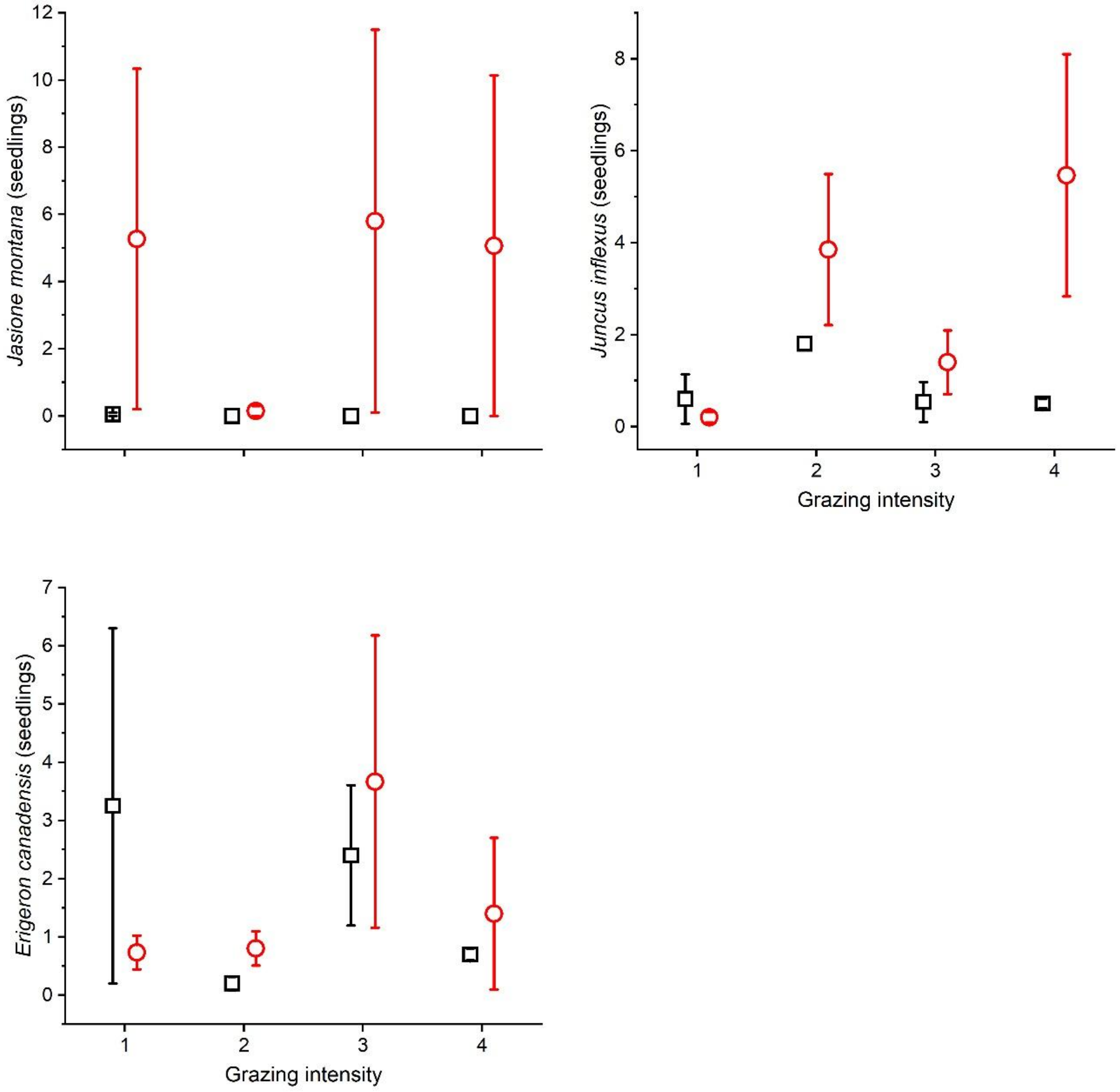
