## Appendix C for "High-intensity sheep grazing impoverishes soil seed banks in sand grasslands"

**Appendix C.** The CSR strategy composition of seed banks of sheep (black squares) and cattle (red circles) grazed grasslands. Intensity levels 1(A), 2 (B), 3 (C) and 4 (D) are shown. For grazing intensity levels please see Table 1.

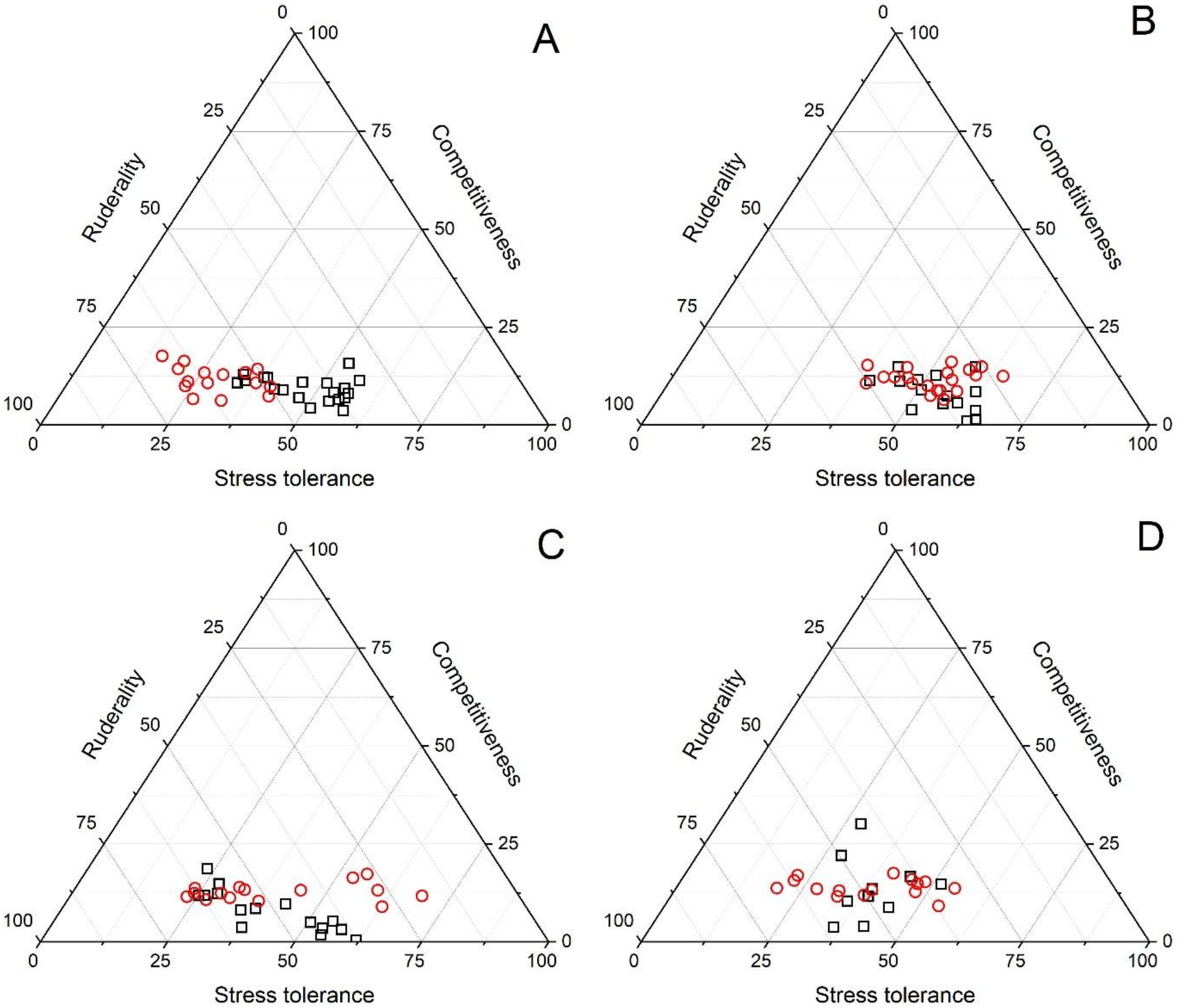
